# Untargeted metabolomics suffers from incomplete data analysis

**DOI:** 10.1101/143818

**Authors:** Richard Baran

## Abstract

**Introduction:** Untargeted metabolomics is a powerful tool for biological discoveries. Significant advances in computational approaches to analyzing the complex raw data have been made, yet it is not clear how exhaustive and reliable are the data analysis results.

**Objectives:** Assessment of the quality of data analysis results in untargeted metabolomics.

**Methods:** Five published untargeted metabolomics studies acquired using instruments from different manufacturers were reanalyzed.

**Results:** Omissions of at least 50 relevant compounds from original results as well as examples of representative mistakes are reported for each study.

**Conclusion:** Incomplete data analysis shows unexplored potential of current and legacy data.

## Introduction

Mass spectrometry-based metabolomics is a powerful tool for the discovery of novel compounds, metabolic capabilities, and biomarkers (Patti el at. 2012; Sévin et al. 2015). Successful discoveries are dependent on the ability to reliably detect relevant signals in raw data and to correctly interpret the underlying spectral features of compounds (Kind & Fiehn 2007; Dunn et al. 2013; Scheubert et al. 2013; Baran & Northen 2013; Kind et al. 2017). The challenging complexity of the data analysis process is well recognized and computational tools facilitating the data analysis process are available (Weber et al. 2017). However, it is not clear how exhaustive and reliable are the current data analysis results. The quality of the results is important not only in the context of exploratory research but even more more so in the context of a strengthening trend towards large scale integration of multi-omic datasets (Perez-Riverol et al. 2017). Public repositories of metabolomics data, such as the UCSD Metabolomics Workbench (Sud et al. 2016) or the MetaboLights (Haug et al. 2013) database, provide an opportunity to reanalyze published raw data to assess the coverage of relevant signals as well as the quality of mass spectra interpretation.

Five untargeted metabolomics datasets from public repositories acquired using instruments from different manufacturers were selected for reanalysis (Table 1, Supplementary Fig. 1-5). The selection was arbitrary with a focus leaning towards high complexity of the raw data (large numbers of detected compounds).

**Table 1.** Untargeted metabolomics studies selected for reanalysis

| Study Identifier | ST000403 | ST000326 | ST000220 | MTBLS214 | ST000259 |
| --- | --- | --- | --- | --- | --- |
| **Instrument** | Thermo Scientific Q-Exactive Orbitrap | Agilent 6530 QTOF | Waters Synapt-G2 Si | AB Sciex TripleTOF 5600 | Bruker MicrOTOF II |
| **Sample Layout** | 6 groups of 3 replicates<sup>a</sup> | 19 individual samples | 3 groups of 7 replicates | 3 groups of 4-5 replicates | 14 groups of 5-6 replicates |
| **Compounds /features (+)** | 590 | 962 | 1259 | 18 | 857 |
| **50+ Omissions** | + | + | + | + | + |
| **Mistakes** | + | + | + | + | + |
| **Study URL** | http://www.metabolomicsworkbench.org/data/DRCCMetadata.php?Mode=Study&StudyID=ST000403 | http://www.metabolomicsworkbench.org/data/DRCCMetadata.php?Mode=Study&StudyID=ST000326 | http://www.metabolomicsworkbench.org/data/DRCCMetadata.php?Mode=Study&StudyID=ST000220 | http://www.ebi.ac.uk/metabolights/MTBLS214 | http://www.metabolomicsworkbench.org/data/DRCCMetadata.php?Mode=Study&StudyID=ST000259 |

<sup>a</sup>Four of the six groups contained added stable isotope labels. Peaks corresponding to clear stable isotope labeling signals or peaks absent from the two control groups without stable isotope labeling were not considered as possible compound omissions.

## Materials and Methods

Raw datafiles along with accompanying data analysis results were downloaded from the respective data repositories (Table 1). Raw data files in original instrument manufacturers’ proprietary data formats were converted to mzXML (Pedrioli et al. 2004) data format using ProteoWizard’s msconvert tool (Chambers et al. 2012). Differences among datasets within a specific study (for ions not reported in original study results) were detected using direct comparisons between datasets binned along the m/z dimension as described previously (Baran et al. 2006). The mass spectra and extracted ion chromatograms corresponding to candidate differences were then inspected visually to assign related ions (e.g. [M+H]^+^, adducts, multimers, in-source fragments, isotopic peaks). To limit the extent of tedious manual curation, the aim of the reanalysis was to find 50 relevant omissions in each study.

To be considered an omission, none of the ions corresponding to the omitted compound could be reported in the original results (even if the only reported ion corresponds to an isotopic peak of an in-source fragment ion of a specific compound). Only raw data acquired in positive mode polarity were used for re-analysis for each study. However, negative mode raw data and results were examined as well. If none of the ions of a specific compound were reported in positive mode results, but at least one ion related to the compound was reported in negative mode results, the compound was not considered and not reported as an omission.

Multiple ions for omitted compounds along with their peak areas are listed in Supplementary Data 1. These lists of ions are not exhaustive. Low intensity isotopic peaks or ions that could be potentially related (but not showing clear similarities in chromatographic profiles, relative peak areas across samples, or differences in *m/z* to other ions of typical chemical relationships) may have been left out of these lists. However, records for even these possibly related ions were sought in original results accompanying the study to make the best effort to report truly omitted compounds in reanalysis results.

Peak areas were calculated using the trapezoidal integration method without any prior smoothing of extracted ion chromatograms or baseline subtraction. Integration bounds were set manually. The ion with the largest peak area from a group of related ions was selected as a “representative” ion for a given compound and used for extracted ion chromatograms (Fig. 1, Supplementary Fig. 6-10). Few representative mistakes found during the reanalysis process were mostly related to ion type (mis)interpretation in the original results and are shown in Supplementary Figures 12-16. A rough comparison of relevance of omitted compounds to the original results was based on peak areas of “representative” ions and a measure of a statistical significance of a difference among the groups of replicate samples in a study, if applicable (Supplementary Fig. 11). Peak areas calculated by the trapezoidal method were normalized to peak areas in the original results (Supplementary Fig. 17-21) for this comparison.

**Figure 1.**
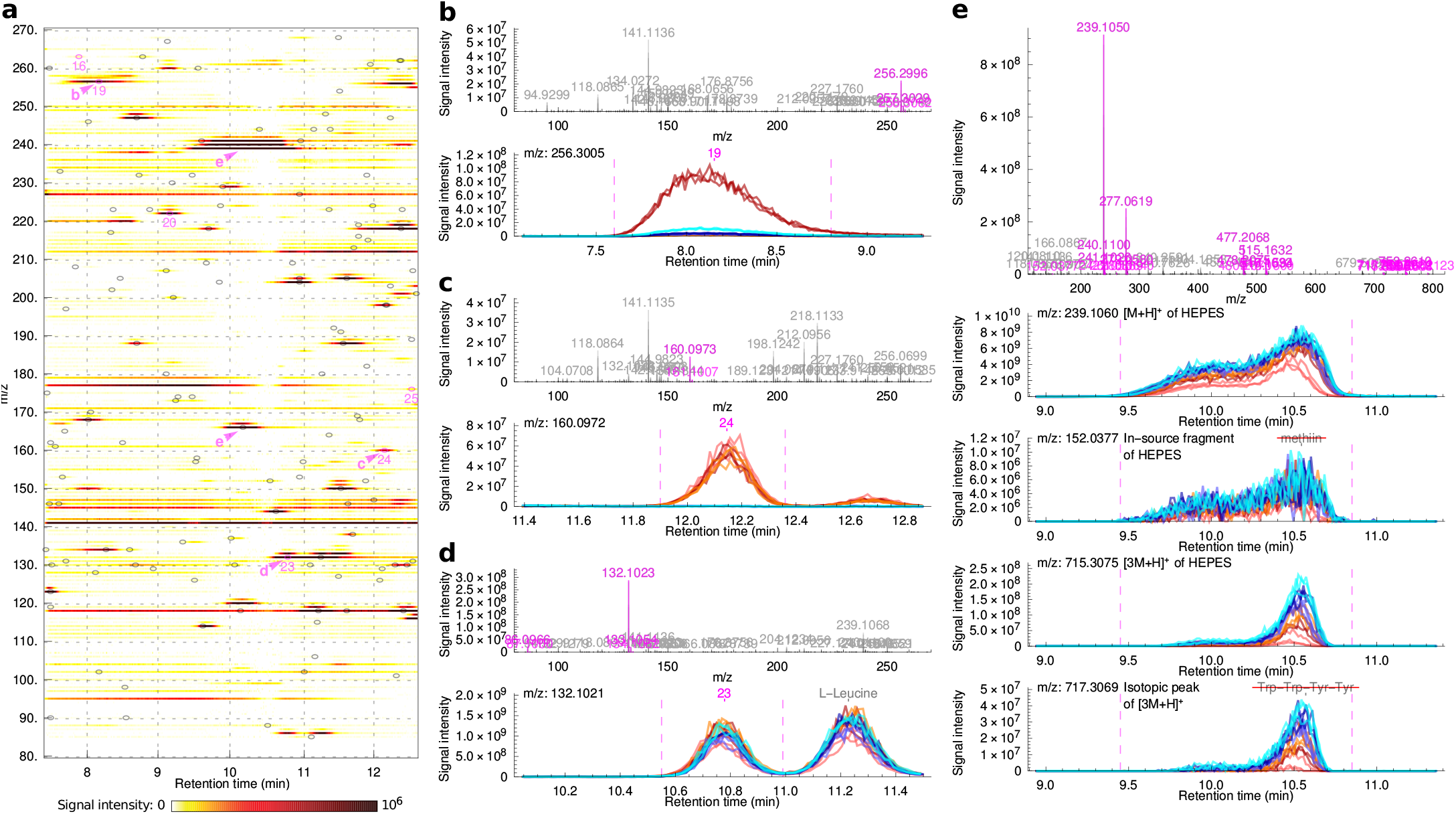
Examples of omissions and mistakes in results from study ST000403. (**a**) Visualization of a part of one of the raw datafiles. Gray labels correspond to annotations from original results accompanying the study data. Magenta labels correspond to omissions or mistakes. (**b-d**) Mass spectra and extracted ion chromatograms for examples of omissions. (**e**) A mass spectrum and extracted ion chromatograms for examples of mistakes. An in-source fragment ion and an isotopic peak of a multimer of HEPES were incorrectly identified as different compounds. Peaks of related ions for a given compound in plots of mass spectra are highlighted in magenta. Color coding for groups of replicate samples in extracted ion chromatograms is the same as in Supplementary Figure 6.

## Results and Discussion

The raw data were reanalyzed as described in the Materials and Methods section to look for omissions of relevant compounds as well as examples of common mistakes in the original data analysis results accompanying the study data. To limit the extent of tedious manual curation of the data, a goal of finding 50 relevant omissions in each study was set. For a compound to be considered omitted, none of its ions (e.g. [M+H]^+^, adducts, multimers, in-source fragments, isotopic peaks) could be reported in the original results. Figure 1a-d shows a few examples of omissions from one of the reanalyzed studies, and Supplementary Figures 6-10 show examples of at least 50 omissions from each study. These omissions are relevant in the context of reported results, since these compounds show either intense signals or differ significantly among the study groups (Supplementary Fig. 11). In addition to omissions, mistakes in ion type interpretation were also found during the reanalysis. The most commonly observed mistake was the reporting of in-source fragment ions, isotopic peaks, or other ion types instead of the protonated molecule [M+H]^+^ ion (Fig. 1e, Supplementary Fig 12-16).

This reanalysis of published metabolomics studies was far from exhaustive. The newly reported lists of ions for omitted compounds (Suppementary Data 1) are incomplete, may contain mistakes as well, and additional unreported compounds are very likely present in the raw data. The selected metabolomics studies have impressive quality of the raw data as well as original data analysis results which must have required significant effort and insight. And yet the results of this simple reanalysis point to an additional unexplored potential of current as well as legacy metabolomics data. Hopefully, these results will strengthen the appreciation for the complexities of the data analysis process and further motivate improvements in computational tools and knowledgebases for metabolomics data analysis.

